# Engineered CRISPR-Cas9 system enables noiseless, fine-tuned and multiplexed repression of bacterial genes

**DOI:** 10.1101/164384

**Authors:** Antoine Vigouroux, Enno Oldewurtel, Lun Cui, Sven van Teeffelen, David Bikard

**Author notes:** Correspondence should be addressed to: Sven van Teeffelen David Bikard.

## Abstract

Over the past few years, tools that make use of the Cas9 nuclease have led to many breakthroughs, including in the control of gene expression. The catalytically dead variant of Cas9 known as dCas9 can be guided by small RNAs to block transcription of target genes, in a strategy also known as CRISPRi. Here, we reveal that the level of complementarity between the guide RNA and the target controls the rate at which dCas9 successfully blocks the RNA polymerase. We use this mechanism to precisely and robustly reduce gene expression by defined relative amounts. We demonstrate broad applicability of this method to the study of genetic regulation and cellular physiology. First, we characterize feedback strength of a model auto-repressor. Second, we study the impact of copy-number variations of cell-wall synthesizing enzymes on cell morphology. Finally, we demonstrate that this system can be multiplexed to obtain any combination of fractional repression of two genes.

## Introduction

A powerful tool to investigate genes and their regulation is to vary their expression level and investigate the response of the cell. Recently, it has been shown that genes can be knocked down from their native locus to varying degrees using CRISPR technology ^1,2^. The RNA-guided Cas9 nuclease from *Streptococcus pyogenes* can be easily reprogrammed to bind any position of interest on the chromosome, with the requirement of an “NGG” protospacer adjacent motif (PAM). The catalytic mutant dCas9 is unable to cleave target DNA, but still binds DNA strongly. It is able to block transcription initiation when binding a promoter region as well as transcription elongation when targeting downstream of a promoter.

Target search begins by probing DNA for the presence of a PAM followed by DNA melting and RNA strand invasion ^3,4^. This last step is extremely sensitive to mutations in the PAM-proximal region known as the seed sequence. Chromatin immunoprecipitation and sequencing assays (Chip-seq) have revealed that several mismatches in the PAM-distal region can be tolerated by dCas9 whose binding can be detected with as little as 5 base pairs (bp) of homology between the seed sequence and the target ^5,6^. These observation are consistent with CRISPRi assays performed in bacteria, which show the ability to reduce transcription of a target gene with as little as 8 bp of homology in the PAM-proximal region ^1^. Accordingly, the degree of gene repression can be controlled quantitatively in two different ways: First, by changing the level of dCas9 expression from an inducible promoter, which then impacts the probability of dCas9 binding to target DNA. This has recently been demonstrated in *Bacillus subtilis* where dCas9 was placed under the control of a xylose-inducible promoter ^7^, as well as in an *E. coli* strain modified to enable tunable control of expression from a P_BAD_ promoter^8^. Second, by introducing mismatches between the guide RNA and the target DNA, as demonstrated in *E. coli* ^1^. While a perfectly matched guide RNA leads to very strong repression, decreasing complementarity in the PAM-distal region progressively reduces the repression strength ^1^.

Here we compare these two repression strategies by characterizing the properties of dCas9-mediated repression at the single-cell level. This enables us to propose a novel physical model of dCas9-mediated repression. It was previously assumed that decreased levels of guide RNA complementarity would decrease repression strength by virtue of reduced occupancy of the target by dCas9 ^9^. Here, we demonstrate a different mechanism: If the target is inside an open reading frame (ORF), complementarity determines the rate at which RNA polymerase (RNAP) kicks out dCas9 during transcription attempts. If dCas9 levels are high enough to saturate the target this mechanism alone determines repression strength. This leads to desirable properties: First, relative repression strength is independent of native expression levels. Second, repression does not add any extrinsic noise to gene expression. On the contrary, tuning gene expression by changing the level of dCas9 expression is inherently noisy and depends on the promoter strength of the target.

We propose to use complementarity-based CRISPR knock-down in combination with fluorescent-protein reporters inserted upstream of a gene of interest to precisely and robustly control its expression. The use of reporter gene fusions rather than direct targeting of the gene of interest yields a predictable repression fold as characterized in this study and provides an easy way to monitor expression levels in single cells. We demonstrate the versatility of our approach using two examples: First, the accurate control of the rate at which the RNAP kicks out dCas9 enables us to quantify the degree of feedback in a model auto-repressor by measuring how much actual gene expression differs from the controlled rate. Second, we take advantage of the ability to obtain a precise degree of repression during steady-state growth to investigate the impact of expression level of an operon coding for two essential cell-wall synthesis enzymes of the ‘rod’ complex, PBP2 and RodA. Finally, we demonstrate that this system can be easily and robustly multiplexed to obtain any combination of the fractional repression of two genes.

## Results

### Varying levels of guide RNA-target complementarity enables noiseless control of gene expression

To quantify how CRISPR/dCas9 modulates gene expression at the single-cell level, we integrated expression cassettes for two constitutively expressed reporters, *sfgfp* coding for the superfolder green fluorescent protein (GFP) and *mCherry* coding for a red fluorescent protein (RFP) at two different chromosomal loci of *E. coli* strain MG1655. To repress either of these genes using CRISPR knock-down we integrated the *dcas9* gene from *S. pyogenes* under a P_tet_ promoter, inducible by the addition of anhydrotetracycline (aTc) ^2^. We then guided the dCas9 protein to target the coding strand of GFP- and RFP-coding ORFs using a constitutively expressed CRISPR array coding for the guide RNAs and the necessary tracrRNA, which form a complex together with dCas9 ^10^.

In this system, repression strength can be tuned in two different ways: either by modulating dCas9 expression level using different aTc concentrations, or by modulating spacer complementarity to the target gene using different numbers of mismatches at the 5’ side of the spacer (Fig. 1, a and b). We employed these two different strategies to repress GFP by different amounts and measured GFP concentration at the single cell level by high-throughput microscopy. As expected, average GFP levels decreased with increasing aTc concentration or increasing spacer complementarity. However, the distributions of single-cell GFP concentrations differed significantly between the two different modes of repression modulation. Specifically, using a perfectly matched guide RNA and varying aTc concentrations led to large cell-to-cell fluctuations, presumably due to variations in dCas9 levels. On the contrary, using high levels of dCas9 expression and varying the degree of guide RNA complementarity maintained the noise (standard deviation over the mean) of single-cell GFP concentration almost constant (Fig. 1, c-d and Supplementary Fig. 1). Noise increased only for very high repression strength, presumably largely due to fluorescence measurement error and possibly due to the stochastic nature of transcription referred to as intrinsic noise, which is only relevant at very low transcription rates ^11^. The plateau value of the expression noise of about 0.3 (corresponding to cell-to-cell variations of 30%) is similar to measurements made by others for constitutive genes in wild-type *E. coli* ^11^. This noise-less repression is qualitatively different from gene repression using transcriptional repressors, which can increase the extrinsic part of the noise of their targets by 5-fold as compared to the unrepressed case ^12^. Accordingly, a similar increase of noise is observed if repression is modulated by inducer concentration (see Supplementary Fig. 1 and also reference ^8^). The alternative system proposed here thus enables to tune expression levels with high precision in single cells.

**Figure 1.**
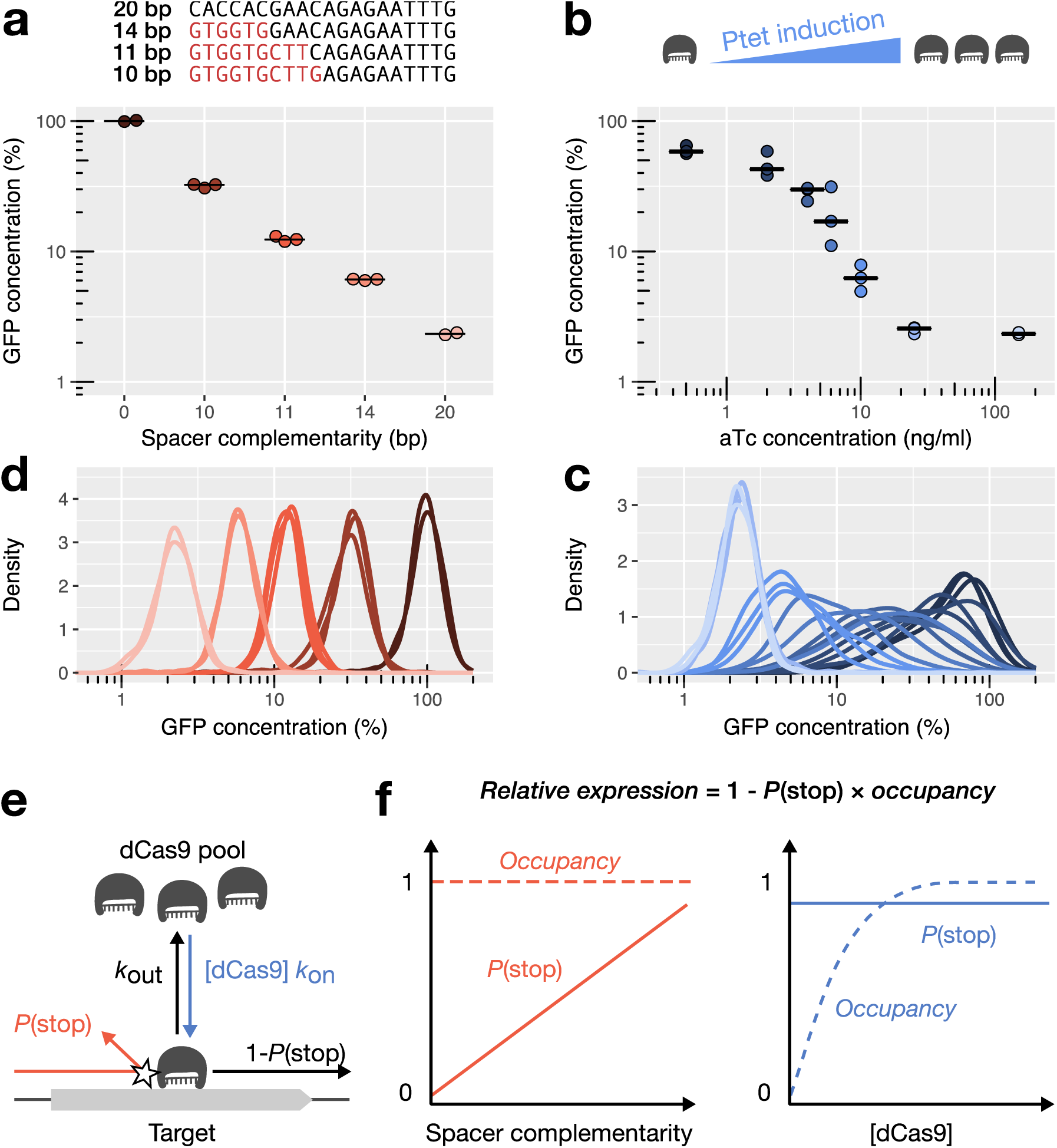
In saturating conditions, CRISPR knock-down can modulate gene expression over a large dynamic range without generating noise. **a, b:** Average cellular GFP concentration obtained (a) by changing guide RNA-target complementarity at a constant high dCas9 concentration or (b) by varying dCas9 levels with increasing concentration of the aTc inducer. Relative GFP concentrations are obtained by high-throughput microscopy and given relatively to the non-targeting spacer at high dCas9 expression. Individual points represent independent replicates. Horizontal bars represent the median of three replicates. **c, d:** Distribution of GFP concentrations for each experiment in panels a and b. Curves of the same color represent replicates of the same condition. **e:** Mechanistic model of dCas9-mediated repression. The expression level of a dCas9-targeted gene is reduced by the product of two probabilities: the probability *P*(stop) of dCas9 blocking RNAP upon collision if occupying the target, and the probability of dCas9 occupying the target (termed occupancy). If dCas9 does not block RNAP during a collision between dCas9 and RNAP, dCas9 is kicked out of the target site (with probability 1-*P*(stop)). The occupancy is determined by binding constant *k*on, dCas9 concentration [dCas9], and dCas9 unbinding rate *k*off. The unbinding rate *k*off, in turn, is the sum of equilibrium unbinding rate and kick-out rate due to collision with the RNAP (see supplements for details). **f:** The two panels schematically illustrate the behavior of the probability of dCas9 blocking RNAP *P*(stop) and dCas9 occupancy if repression strength is controlled by guide-RNA complementarity (left) or dCas9 concentration (right), respectively.

To explain these observations, we formulate a model of dCas9 repression according to which, for a high enough intracellular concentration of dCas9, the target becomes saturated by dCas9 but transcription may still occur when dCas9 is kicked out by the RNAP. In this model, mismatches affect the repression level by increasing the probability that dCas9 is ejected upon collision (see Supplementary Information). Our model thus makes the prediction that gene repression should be independent of dCas9 concentration even in the presence of mismatches between the guide RNA and the target, as long as there are enough dCas9 complexes to saturate the target. We could indeed verify this prediction by reducing the fraction of active dCas9 complexes using a decoy guide RNA (Supplementary Fig. 2). When repression strength is controlled by mismatches rather than dCas9 concentration, the cell-to-cell fluctuations of dCas9 concentration no longer affect the repression of the target, thus explaining the low noise obtained in Figure 1c and Supplementary Figure 1

**Figure 2.**
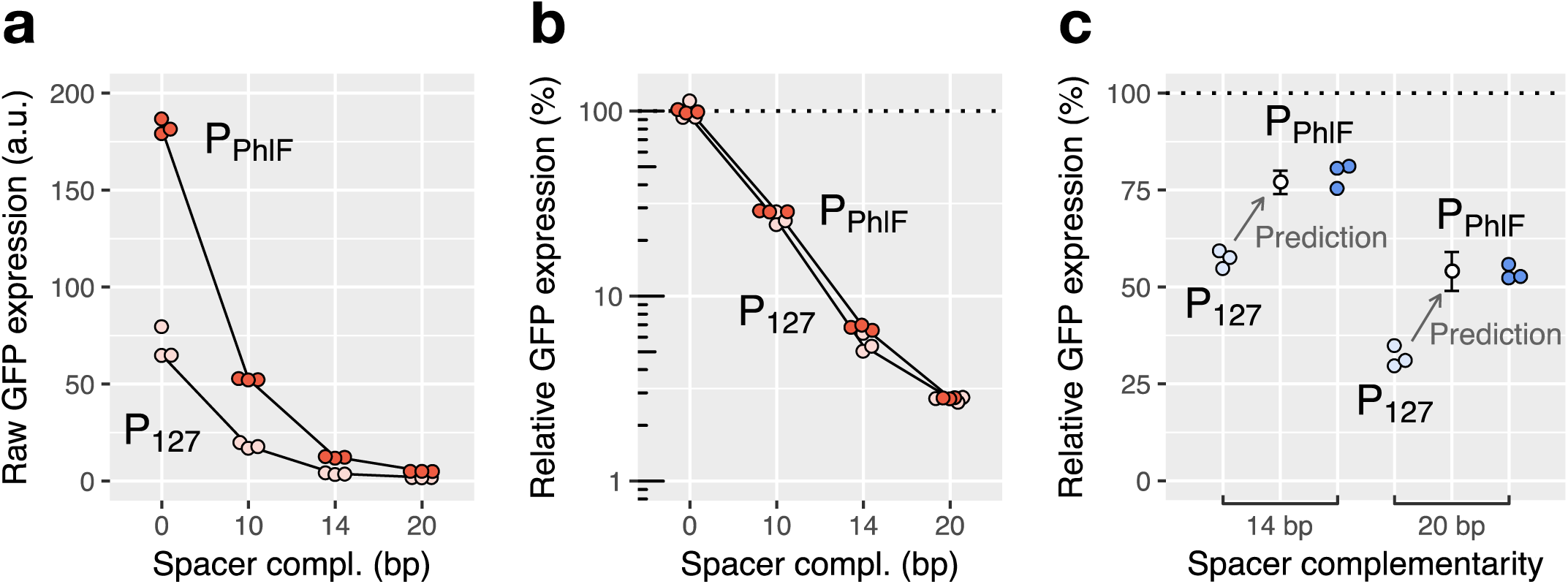
Relative repression by dCas9 is independent of promoter strength only in saturating conditions. Relative GFP expression measured by flow cytometry for two promoters of different strengths (P127 and PPhlf) and repressed using the same set of spacers for saturating (a, b) and non-saturating (c) dCas9 concentrations. **a, b**: Raw GFP expression (a) and relative GFP expression with respect to a non-targeting spacer (b) for a saturating dCas9 concentration. While PPhlF is about 3 times stronger than P127, the relative expression levels after repression are similar for both promoters. **c:** Experimental and predicted relative GFP expression for a non-saturating dCas9 concentration (using a 40-times lower concentration of aTc). Repression is weaker for the stronger PPhlf promoter for up to 6 mismatches on the guide RNA, in quantitative agreement with the kick-out model (see Supplementary Information). Error bars: standard error of the mean of the computational prediction‥

### The relative repression level is robust with respect to changes in the target’s promoter strength

To use CRISPR knock-down on genes with different native expression levels, it is important to know whether the transcription rate of the target has an influence on the relative repression. Our model predicts that repression strength should not be measurably affected for promoters of different strengths, if dCas9 is saturating the target as already indicated above. To verify this prediction, we put *sfgfp* under the control of two promoters of different strengths (P_127_ and P_Phlf_) and blocked expression using four different guide RNAs with an increasing number of mismatches. While the strain with P_PhlF_ expressed about three times more GFP than the strain with P_127_ (Fig. 2a), the repression fold with regard to the promoter’s initial expression level was identical in each case (Fig. 2b). These observations confirm that repression by mismatched guide RNAs in saturating conditions is independent of promoter strength. On the contrary, if repression is controlled by varying dCas9 concentration, our model predicts that repression fold is dependent on promoter strength: a stronger promoter is expected to kick out dCas9 more frequently, reducing the occupancy of the target. Indeed, we observed a weaker repression of P_Phlf_ compared to P_127_ at low dCas9 concentrations (Fig. 2c) that quantitatively agrees with our model prediction (Fig. 2c and Supplementary Information).

### dCas9 ejection probability increases with temperature

It was recently reported ^13^ that dCas9 is no longer active at 42°C, suggesting that repression strength might decrease with increasing temperature. This observation also bears the possibility that our system becomes less robust with respect to dCas9-copy number fluctuations and promoter strength with increasing temperature, if the condition of target saturation was not fulfilled. To quantify the temperature dependence of repression and test for robustness, we measured the repression of RFP by guide RNAs with 11 bp or 20 bp of complementarity at temperatures ranging from 30 to 42°C. The repression strength decayed continuously with increasing temperature (Fig. 3), displaying a sharp decrease of repression between 37 and 42°C. Regardless of the temperature, repression strength was not affected by dCas9 complex concentration (Supplementary Fig. 3). From our model we can thus conclude that increasing temperature does not affect dCas9 occupancy but increases the probability of dCas9 being kicked out by the RNAP. This also indicates that our system should work independently of promoter strength at all temperatures tested.

**Figure 3.**
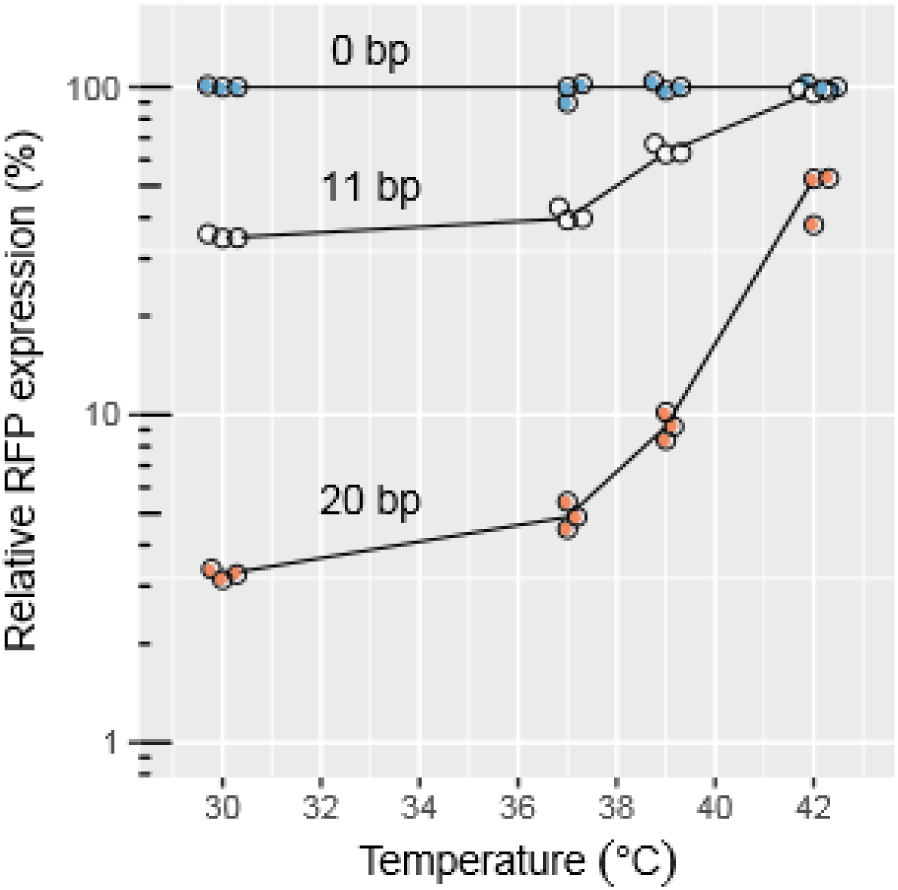
The efficiency of CRISPR knock-down is affected by high temperatures. Relative RFP expression measured by flow cytometry upon repression with different levels of complementarity and at different temperatures. The values are normalized with respect to the non-targeting spacer at each temperature.

### CRISPR knock-down in combination with fluorescent-protein insertions can be used to repress and monitor genes in their native contexts

Precision, robustness, and large dynamic range make complementarity-based CRISPR knock-down a versatile repression strategy. To repress genes of interest in their native context, we propose to insert *sfgfp* or *mCherry* reporters as transcriptional or translational fusions upstream of the gene. A library of CRISPR plasmids can then be introduced to repress the fusions to the desired levels by targeting the *sfgfp* or *mCherry* coding sequences. The method thus allows taking advantage of the measured repression levels for constitutive promoters established above. Furthermore, gene expression can be measured at the single-cell level, revealing cell-to-cell variations. In the following, we demonstrate that this system has broad applicability for the study of genetic regulation and cellular physiology: first, we study the regulation of a model transcriptional feedback circuit, and second we quantify the effect of fractional protein repression on cell morphology during steady-state growth.

### CRISPR knock-down can be used to uncover and characterize genetic feedback

To demonstrate the versatility of our system for the study of genetic circuits, we chose the previously described PhlF auto-repressor from *Pseudomonas fluorescens*15 as a model system: We constructed a synthetic operon consisting in the P_PhlF_ promoter followed by the *sfgfp* and *phlF* genes in a single operon (Fig. 4a). The PhlF repressor binds to the P_PhlF_ promoter and decreases transcription initiation, thus creating an artificial negative feedback loop. The strength of this feedback can be externally reduced by adding the chemical inducer 2,4-diacetyl-phloroglucinol (DAPG) that blocks binding of PhlF to the promoter (Supplementary Fig. 5). Accordingly, higher DAPG concentrations lead to higher steady-state concentrations of PhlF and GFP. To quantify the feedback strength for different DAPG concentrations we targeted the *sfgfp* ORF using spacers with variable degrees of complementarity (Fig. 4). CRISPR knock-down of GFP should lead to an increased transcription initiation rate of the promoter. As a consequence the fold change of expression during CRISPR knock-down should be lower in the case of feedback than without feedback. The quantitative difference between the two situations can then be used to quantify the feedback strength.

**Figure 4.**
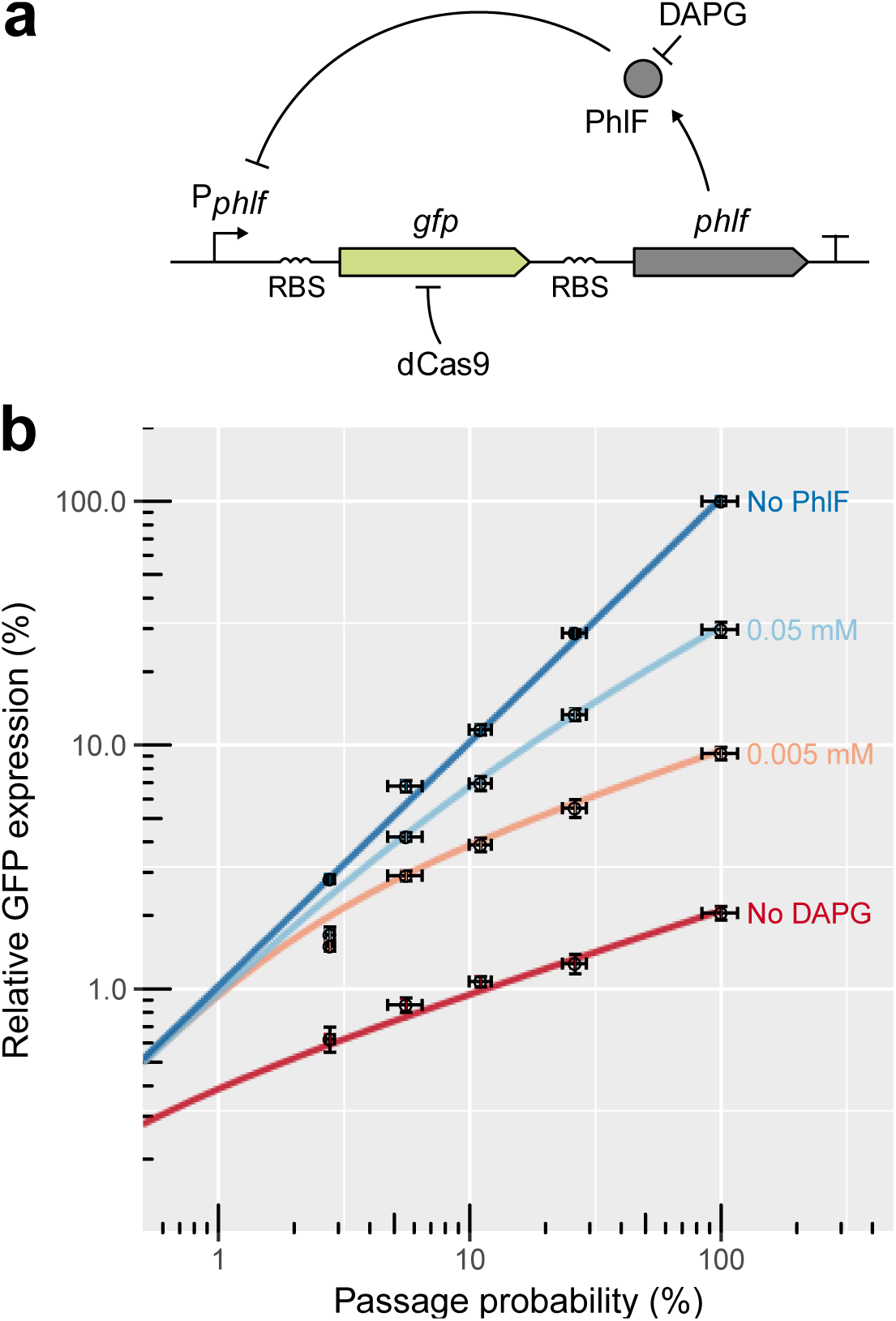
CRISPR knock-down can be used to quantitatively characterize feedback loops. **a:** Schematic of the synthetic feedback loop constructed for this experiment. The strength of the feedback can be modulated by addition of DAPG, an inhibitor of PhlF. RBS: Ribosome Binding Site. T: Transcription terminator. **b:** Flow cytometry measurements and fits to a theoretical model of relative GFP expression levels, where GFP is expressed from the artificial feedback loop presented in panel a. GFP expression is normalized by the maximal level of GFP expressed constitutively from the PPhlF promoter alone (indicated as “No feedback”). The GFP is repressed using 4 different guide RNAs with respectively 10, 11, 14 and 20 bp of complementarity. The passage probability 1-*P*(stop) associated with each of these guide RNAs was measured in parallel on a strain expressing GFP constitutively from the P127 promoter. Adding different amounts of DAPG to the medium reduces the strength of the feedback, causing the steady-state level to increase and repression to become more efficient. The colored lines represent the GFP expression as predicted by a mathematical model that was fitted to the data (see Supplementary Information). For each DAPG concentration, a binding constant characterizing the strength of the feedback and a Hill coefficient were determined. Error bars: 95% confidence interval of the mean based on 3 biological replicates.

**Figure 5.**
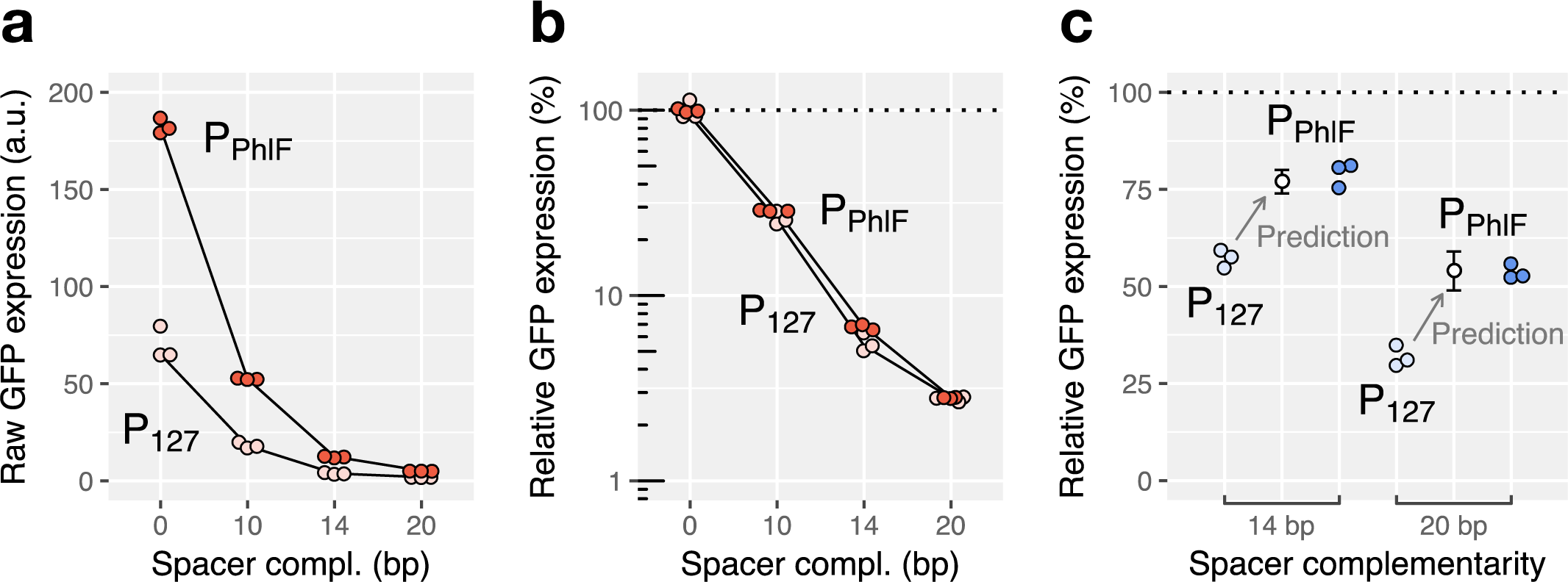
CRISPR knock-down of the *mrdAB* operon increases cell width at high repression strengths. **a:** Schematic of the modified chromosomal locus of the *mrdAB* operon in strain AV08. **b:** Cell shapes observed by phase-contrast microscopy for cells grown in M63 minimal medium. Different repression levels of the *mrdAB* operon are compared to wild-type *E. coli*. Cells with different cell lengths were picked at random and images were rotated numerically. **c:** Cell width as a function of the mCherry-PBP2 concentration measured by fluorescence microscopy. Each point represents a cell and colors represent different levels of spacer complementarity. The connected white dots represent the population averages (mean of 3 biological replicates). The dotted line represents the average cell width for wild-type *E. coli* (mean of 3 replicates). The values are normalized with respect to the non-targeting spacer. **d, e:** Linear regression between mCherry-PBP2 concentration level and cell width, for the strains repressed with 11 bp (panel d) and 0 bp (no repression, panel e). rS is the Spearman correlation coefficient (median of 3 biological replicates). The negative value indicates that cells with a lower level of PBP2/RodA tend to be wider. The p-values (two-sided F-test) measure the certainty that the slope is different from 0.

As anticipated, expression of GFP decreased with increasing complementarity and the relative reduction of expression was less pronounced with feedback than without feedback (Fig. 4b). We then fit the expression data to a mathematical model of gene repression (Fig. 4b and Supplementary Information) to calculate for each DAPG concentration the binding constant of the repressor *K* and a Hill coefficient *n*. We observe that a Hill coefficient of *n*=2 describes our data for low DAPG concentrations (0 μM and 5 μM), while a Hill coefficient of *n*=1 was required to describe our observations at 50 μM. PhlF proteins dimerize *in vitro* and are thought to bind the operator as a dimer ^16^. We thus hypothesized that PhlF is predominantly found as monomers at low DAPG concentrations, where PhlF expression is also low, and as dimers at high DAPG concentrations, where PhlF expression is high. The role of DAPG concentration in this transition is discussed in the Supplementary Information.

The detailed insights obtained here demonstrate the usefulness of precisely controlling the rate at which the RNAP is blocked by dCas9, while monitoring residual expression with a fluorescent reporter. The same method can be applied to other and more complex problems of gene regulation, e.g., by monitoring the response of one gene to the precisely tuned levels of another gene repressed by CRISPR knock-down.

### CRISPR knock-down reveals how cells adapt their shapes to low levels of an essential cell-wall synthesis operon

We then used our approach to explore the morphological response of cells to different expression levels of two essential proteins for peptidoglycan cell-wall synthesis encoded by the *mrdAB* operon. PBP2 (encoded by *mrdA*) and RodA (encoded by *mrdB*) are inner membrane proteins with respectively transglycosylase ^17^ and transpeptidase activity ^18^. The two highly conserved enzymes are part of the multi-enzyme ‘rod’ complex, which is essential for cell-wall synthesis during cell elongation ^19^.

Previous depletion experiments suggest that PBP2 expression is buffered against large copy-number fluctuations, as cells grow for multiple generations before showing a reduction of growth rate ^20^. The drawback of depletion experiments is that they do not allow studying the effect of protein abundance in the steady state. To quantify the relation between PBP2 levels and morphological response during steady-state growth we constructed a translational protein fusion by seamlessly integrating *mCherry* in front of the *mrdA* open reading frame in the native chromosomal *mrdAB* locus (Fig. 5a).

The mCherry-PBP2 fusion is fully functional, similarly to a fusion constructed previously^20^. We then introduced a chromosomal P_tet_-*dCas9* cassette and different pCRRNA plasmids programmed to target *mCherry* with 0, 11, 18 or 20 bp of complementarity in order to obtain a range of transcription rates for the operon. These strains were induced for dCas9 expression and grown until protein levels and cell dimensions reached steady state (Supplementary Fig. 6). Lowering expression of the *mrdAB* operon led to increasing cell width, with a sharp rise of cell width below ∼20% of the native expression level (Fig. 5b, 5c and Supplementary Fig. 7a), while cell length was largely unaffected except for the highest repression strength (Supplementary Figs. 8b and 9). Cells presumably increased their diameters to lower their surface-to-volume ratio in response to reduced levels of cell-wall synthesis ^21^. In support of this hypothesis, cell-to-cell fluctuations in the intracellular density of mCherry-PBP2 negatively correlated with cell diameter for intermediate *mrdAB* repression (at 11 bp guide RNA/target complementarity) (Fig. 5d). Such a correlation was not observed when the operon was not repressed (Fig. 5e), indicating that the cells buffer natural fluctuations of *mrdAB* and thus avoid fluctuations of cell morphology, as previously suggested ^20^. By gradually lowering the levels of PBP2 and RodA, we were able to take the cells out of the buffering regime at about 30% of native expression. Together, these experiments demonstrate that cells buffer gene expression of an essential operon against copy-number fluctuations of about threefold and that cells cope with even stronger fluctuations by adjusting their surface-to-volume ratio. However, once expression levels are reduced by more than fivefold, cells show severe growth defects.

**Figure 6.**
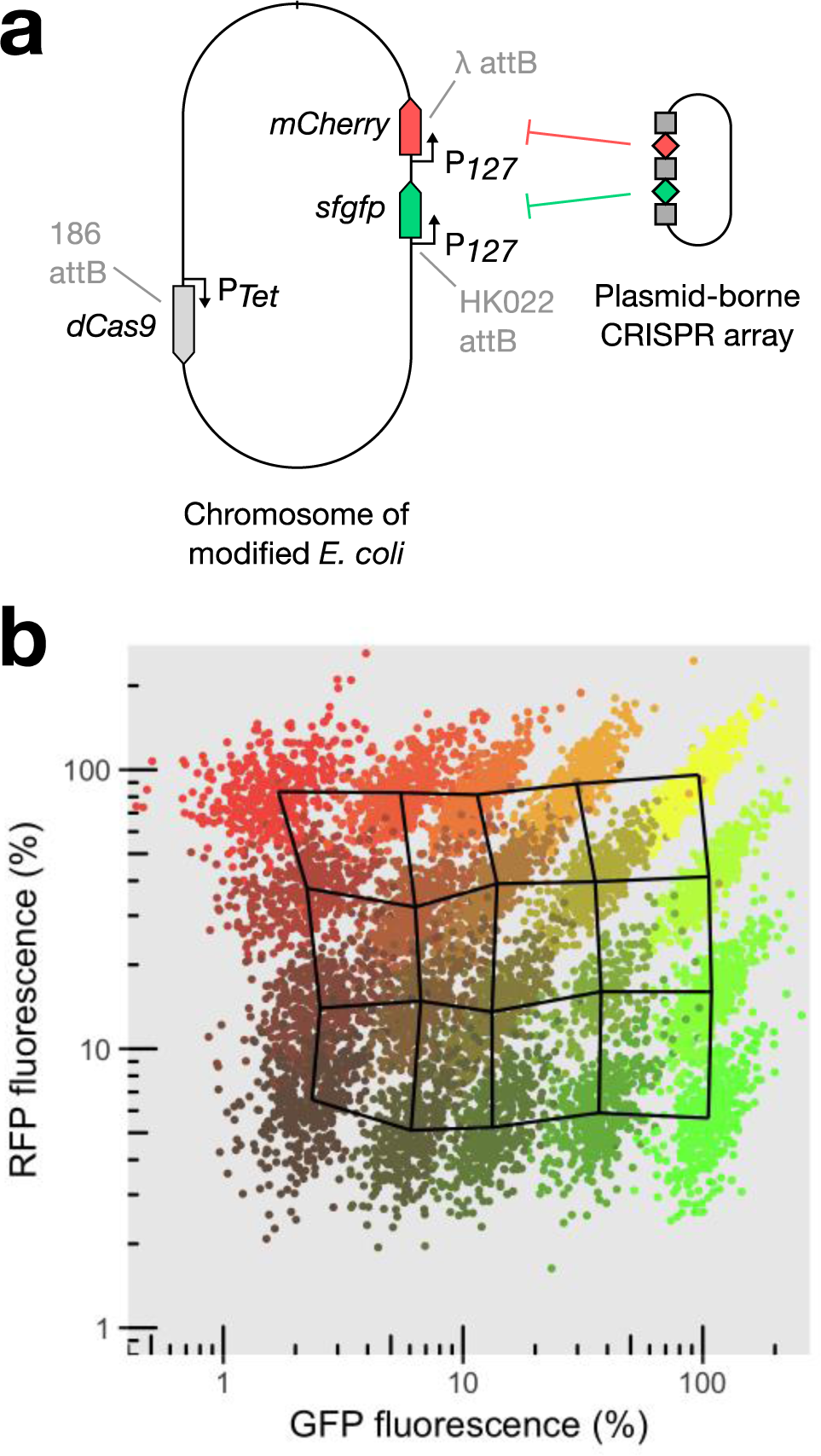
CRISPR knock-down can be multiplexed to modulate expression of two genes without cross-talk. **a:** Schematic of the strain expressing two reporters and PTet-*dCas9* integrated in the chromosome at phage attachment sites. The levels of the two reporters can be controlled using a plasmid-borne CRISPR array coding for guide RNAs (diamonds) interspaced with CRISPR repeat motifs (squares), and also carrying the tracrRNA sequence (not shown). **b:** Relative GFP and RFP concentration given relatively to the non-targeting spacer measured by high-throughput microscopy for a collection of 20 CRISPR plasmids. Each point represents a single cell, and each color represents the population obtained with one CRISPR plasmid. The overlaid meshwork connects the median values of the different populations.

### CRISPR knock-down can be used to modulate the expression of two genes

With our method, the fractional repression level of any target gene is controlled genetically rather than chemically by the concentration of an inducer. It can thus be used to modify expression of multiple genes independently. To assess this potential, we built a library of CRISPR arrays containing two spacers, one targeting *sfgfp* and the other *mCherry*. We selected five spacers with varying levels of complementarity to each of the target, spanning a large range of expression, from 2% to 100% of the initial level. We combined these spacers to form 20 CRISPR arrays that cover the entire space of expression and used them to control the concentrations of GFP and RFP expressed from the chromosome (Fig. 6a). As expected, the repression of one gene is independent of the repression of the other (Fig. 6b). Strong correlations between GFP and RFP in single cells targeted with the same combinations of guide RNAs are due to common sources of extrinsic noise^12^. We anticipate this to be a useful tool to study interactions of genes and specifically the effect of stoichiometry in genetic networks.

## Discussion

Here, we demonstrate that tuning gene expression through complementarity between guide RNA and target works robustly at the single-cell level, with two specific advantages over previous methods relying on inducible promoters: First, relative repression strength is independent of native expression levels, making the system applicable to study genes of vastly different promoter strengths. Second, the system preserves endogenous expression noise of the repressed gene. This allows studying the impact of gene repression on cellular physiology without generating stochastic cell-to-cell variability, which is known to have important downstream consequences for processes such as cell differentiation or the emergence of spatial structure in populations^22,23^.

The ability to control gene expression level through guide RNA complementarity rather than the concentration of an inducer has other advantages: For example, it enables differential control of different cells within the same culture. This could prove useful in pooled screens or competition assays. Furthermore, the strategy can be multiplexed to enable the simultaneous control of multiple genes independently without requiring multiple chemical inducers. We demonstrated this ability with two targets (GFP and RFP; Fig. 6), which can be inserted in front of genes of interest. The strategy can easily be extended to include more than two fluorescent reporters as targets.

Alternatively to using fluorescent-protein fusions it is also possible to guide dCas9 directly to the gene of interest, but this comes with the disadvantage of uncertainty about the exact repression strength due to two reasons: First, the rate at which dCas9 blocks the RNAP is dependent on the specific target sequence. In the future it might thus be desirable to develop computational means to predict target repression based on sequence alone^24^. Second, any feedback controlling the expression of the target could lead to altered transcription-initiation rates (Fig. 4). Therefore, using fluorescent-protein fusions has the advantage to report the exact expression level.

The properties of dCas9 repression described in this study originate from the mechanism of dCas9 binding to DNA inside ORFs. We show that at high concentrations, dCas9 is saturating the target site even when using guide RNAs with large numbers of mismatches, where repression of the target gene is weak. We explain these observations by a ′kick-out′ model of repression, according to which RNAP kicks out dCas9 with a probability that can be tuned by spacer complementarity (Supplementary information). The model predicts that repression is promoter-strength dependent in non-saturating conditions, in quantitative agreement with experiments (Fig. 2c). Our model also predicts a low rate of spontaneous unbinding of dCas9 from the target for full and intermediate (14 bp) complementarity (Supplementary Fig. 10). For full complementarity, our model is compatible with the hypothesis that dCas9 never leaves the target spontaneously but gets kicked out either by the RNAP or during DNA replication. This prediction is consistent with the long half-life of dCas9 binding *in vitro* previously reported^3^. A recent high-throughput study of dCas9 off-target binding and unbinding suggests that mutations in the PAM-distal region control the unbinding kinetics of dCas9^24^. Since unbinding is dominated by RNAP-dCas9 collisions for the promoters tested here and as rebinding to the target is fast at high dCas9 concentrations, spontaneous unbinding plays no significant role for gene repression in our model system. However, if dCas9 targets the promoter rather than the coding region, spontaneous unbinding might be the only mode allowing residual gene expression. This view is supported by the higher repression strength observed when targeting the promoter region^1,2^.

Our results also suggest that dCas9 ejection does not lead to bursts of transcription, but that instead dCas9 returns to the target site after ejection in a time that is small with respect to the typical time interval between transcription initiations. It is still conceivable that such bursts may occur for transcription rates higher than the strongest promoter we used.

Our strategy enables to precisely control gene expression without introducing cell-to-cell variability, and should be useful for any quantitative measurements that depend on the expression level of a gene. By taking advantage of the ability to precisely control the rate at which dCas9 blocks the RNAP we could characterize a synthetic feedback loop, revealing unexpected properties of Phlf repression activity. In a second example, we took advantage of the ability to fine-tune expression levels at the steady state to quantitatively measure cell shape as a function of the levels of PBP2 and RodA. The level of precision achieved here would be hard to establish with conventional methods. Accordingly, this is the first study to establish a quantitative relationship between the abundance of cell-wall synthesizing proteins and cell morphology at the population and single-cell levels. We anticipate that our method will be useful to study many other systems and in particular genetic circuits that include high levels of noise, such as stochastic switches, or other noise-dependent processes, where preservation of a well-defined level of expression noise is desirable.

## Materials and methods

### Genome modifications

All the strains used for measurements derive from *E. coli* MG1655. Table S1 details the construction of the strains used in this study.

For integration of cassettes at phage attachment sites, we used the “clonetegration” method^25^. Integrated backbones were excised by expressing a flippase from pE-FLP. Plasmid pLC97 (addgene number: to be announced) can be used to easily integrate P_Tet_-*dCas9* at the lambda attB site.

For scarless integration of *mCherry-mrdA* in the native *mrdAB* operon we used the pCas / pTarget system^26^. The PAmCherry-PBP2 protein fusion present in strain TKL130^27,20^ was replaced by a translational fusion with *mCherry* extracted from plasmid pFB262^28^. To this end the pAV06 variant of pTarget was constructed and genome editing was performed as described in reference^26^.

The sequences of the *sfgfp* and *mCherry* genes used in this study can be found on the GenBank database with accession codes KT192141.2 and JX155246.1, respectively.

### Plasmid design and construction

The CRISPR targets were chosen next to the beginning of the ORF, but at least 50 bp away from the initiation codon, in order to preclude unwanted interactions with the native regulation of transcription. None of spacers used in this study have any off-target position with more than 8 bp of complementarity in the PAM-proximal region. Spacers were cloned into the CRISPR array of plasmid pCRRNA using Golden-Gate assembly as previously described^29,30^. The oligonucleotide sequences are available in table S4.

The other plasmids from this study were constructed by Gibson assembly^31^. The fragments are described in table S2 and the primer sequences in table S3. DNA constructions were electroporated in *E. coli* strain DH5α or Pi1 for *pir*-dependent origins of replication^32^.

### Media and reagents

For all flow cytometry measurements, the cells were grown in Luria-Bertani (LB) broth. As a minimal medium for the *mrdAB* measurements, we used M63 medium supplemented with 2 g/L of glucose, 10 mg/L of thiamine, 10 mM of MgSO_4_ and 1 g/L of casaminoacids. When needed, we used various antibiotics (25 μg/ml chloramphenicol, 100 μg/ml carbenicillin, 50 μg/ml kanamycin, 100 μg/ml spectinomycin). Di-acetyl-phloroglucinol (DAPG) and anhydrotetracycline (aTc) were used respectively for induction of P_PhlF_ and P_Tet_ promoters. All oligonucleotides were obtained from Eurofins Genomics.

### Preparation of steady-state exponential cultures

Unless stated otherwise, all cultures were grown at 30°C. Strains were first restreaked from a freezer stock. Independent single colonies were picked for each replicate. Cells were then grown overnight in 96 deep-wells plates using a table-top shaker in 1 ml of medium with 100 ng/ml of aTc and 50 μg/ml of kanamycin (Eppendorf). The day of the measurement, cultures were back-diluted 250 times in fresh medium with aTc and kanamycin, and grown for 1h45 into exponential phase. We then fixed the cells with 4% formaldehyde (30 min on ice) and washed with PBS.

### Flow cytometry

Fluorescence of single cells was recorded using a Miltenyi MACSquant flux cytometer. 10,000 events were recorded per replicate. In all cases, the AV01 strain (with no reporters) carrying a non-targeting pCRRNA plasmid was used to measure the auto-fluorescence background. We calculated the mean fluorescence signal of each population and subtracted the mean autofluorescence signal. To test whether differences in expression were significant we performed Student’s t-test on the natural logarithms of the average fluorescence^33^.

### High throughput microscopy (imaging cytometry)

An Amnis ImageStreamX (EMD Millipore) imaging cytometer was used to image the cells in high throughput in brightfield, GFP, and RFP channels. Images were analyzed using the IDEAS® (EMD Millipore) software suite. For each condition, at least 10,000 events were recorded per replicate. Cells that were out of focus or tilted were identified by calculating the average gradient of a pixel normalized for variations in intensity levels (*Gradient RMS* feature in IDEAS®). Additionally, we used the Feature Finder script of IDEAS® to remove contaminating particles, images with multiple cells and beads. After filtering, at least 2000 images remained per sample. The fluorescence channels were not used for filtering. A color compensation matrix was calculated to account for spectral overlap of GFP and RFP emission spectra, so cultures of AV02 (GFP only) and AV04 (RFP only) would each have a null signal on the converse channel. As a proxy for the reporter’s intracellular concentration we used the average image intensity inside the area corresponding to each cell. The cell area was determined by using a threshold on the bright field images. Single points located more than 3 standard deviations away from the population average were discarded as outliers, as they can disrupt the noise computations. The average fluorescence *μ* of each sample was calculated by taking the mean of the single-cell fluorescence. The noise was defined as *σ/μ*, with 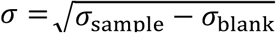 where *σ*_sample_ is the standard deviation of the intracellular average intensity of the sample, and *σ*_blank_ is the standard deviation of a sample with no fluorescent reporter (noise from auto-fluorescence).

### Fluorescence and phase-contrast microscopy

Fixed cells were transferred to Phosphate Buffered Saline (PBS) microscopy pads with 1.5% agar and imaged using an inverted microscope (TI-E, Nikon Inc.) equipped with a 100x phase contrast objective (CFI PlanApo LambdaDM100x 1.4NA, Nikon Inc.), a solid-state light source (Spectra X, Lumencor Inc.), a multiband dichroic (69002bs, Chroma Technology Corp.). mCherry fluorescence was measured using excitation (560/32) and emission (632/60) filters. Images were acquired using a sCMOS camera (Orca Flash 4.0, Hamamatsu) with an effective pixel size of 65 nm.

MatLab code adapted from the Morphometrics package^34^ was used to find cell contours from phase-contrast images. Background intensity, uneven illumination and cell auto-fluorescence were accounted for in the analysis. For the analysis of fluorescence signal, we corrected the raw mCherry values for uneven illumination, background intensity and cell auto-fluorescence. Intracellular protein concentration was obtained as the mean pixel intensity inside the cell area. Total regression was used to find the major axis of the cell. Cell width was defined as the average distance between the cell contour and this axis, excluding the poles.

Measurements of cell morphology during steady-state exponential growth (Supplementary Fig. 6) were performed after overnight induction of dCas9, followed by 1/250 dilution of the culture in fresh M63 medium with aTc and while the culture was kept in exponential growth phase at an optical density below 0.1. Samples were taken from the culture, fixed and imaged after 2h and 4h.

### Northern blot

Total RNA was extracted from cultures in early stationary phase using Trizol. Electrophoresis on Novex® TBE-Urea Gels (10% polyacrylamide gels containing 7 M urea, Invitrogen) was used to separate RNAs. The gels were blotted onto Nylon membranes (Invitrogen), which were subsequently cross-linked with 1-ethyl-3-(3-dimethylaminopropyl) carbodiimide (EDC, Thermo Scientific) buffer^35^. The probes were labeled as follow: 100 pmol of oligonucleotide were heat denatured, labeled and phosphorylated by mixing 40 μCi of 32P-γ-ATP (PerkinElmer) and T4 PNK (NEB) reagents. A labeled probe specific to the guide RNA R20 (5’ GCATAGCTCTAAAACTCCGTATGAAGGCACCCAGA 3’) was column purified (Macherey-Nagel PCR cleanup kit) and used for overnight hybridization. The intensity of the shortest band, corresponding to the fully processed guide RNA, was quantified using the Fiji software package.

## Acknowledgements

We thank T. K. Lee and K.C. Huang for providing strain TKL130, F. Bendezù for providing plasmid pFB262, H. Cho and T. Bernhardt for providing plasmid pHC942 and D. Mazel for providing plasmid pSW23t. We thank J. Fernández-Rodríguez for providing the P_Phlf_-*sfgfp-phlF* DNA fragment. We thank E. Brambilla and E. Oldewurtel for support in microscopy, as well as A. Soler and M. Hasan for support regarding flux cytometry and imaging cytometry. We acknowledge the Technology Core of the Center for Translational Science (CRT) at Institut Pasteur for support in conducting this study. This work was supported by the European Research Council (ERC) under the Europe Union’s Horizon 2020 research and innovation program (grant agreement No [677823] to DB and No [679980] to SvT); the French Government’s Investissement d’Avenir program Laboratoire d’Excellence ‘Integrative Biology of Emerging Infectious Diseases’ [ANR-10-LABX-62-IBEID] to SvT, DB, and AV; the Marie de Paris “Emergence(s)” program to SvT; and the Pasteur-Weizmann consortium to DB.

## Author contributions

DB, SvT and AV designed the study. AV constructed the strains and plasmids and performed all measurements and data analysis. SvT developed the mathematical models. LC took part in strain construction and EO performed the compensation of uneven illumination for the measurement of mCherry-PBP2 by microscopy. AV, DB and SvT wrote the manuscript.

## Conflict of interest

The authors declare that they have no conflict of interest.

